## Supplementary material for "Dissociable laminar profiles of concurrent bottom-up and top-down modulation in the human visual cortex"

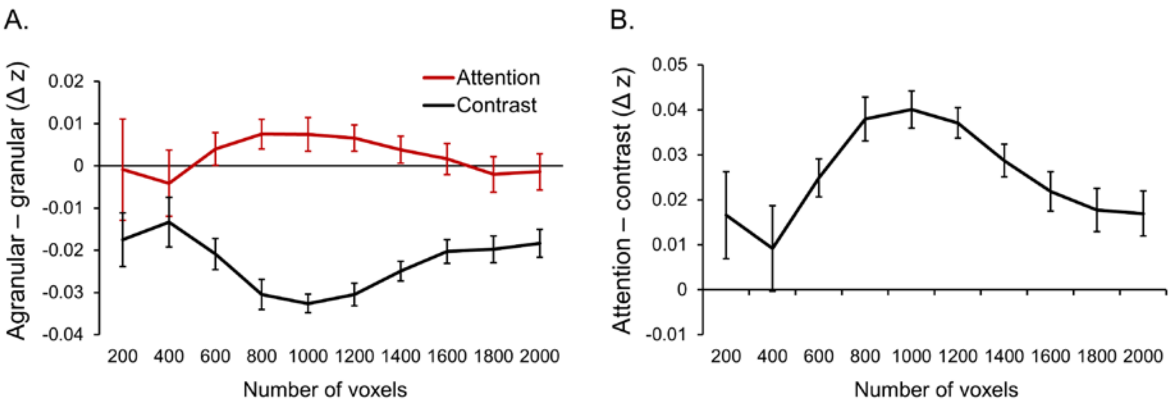


***Figure S1:*** **Effect of mask size on results.** In our main analysis, we constructed ROIs for V1, V2 and V3 comprising the 1000 most selective voxels (500 that preferred clockwise and 500 that preferred counter-clockwise). Here we conducted a series of control analyses to ensure our choice of 1000 voxels did not bias our results. **A:** Agranular-granular scores for the effects of feature-based attention (red) and stimulus contrast (black) for a range of different mask sizes. **B:** Difference between agranular-granular scores for the effects of attention and contrast plotted in panel A. A positive difference indicates that the effect of attention was more agranular compared to the effect of contrast. Though effect sizes varied with mask size, being smaller at the extremes (likely due to increased noise in laminar estimates when using few voxels and the inclusion of less selective voxels when using a large number of voxels), attention was more agranular than contrast for all mask sizes, as was the case in our main analysis. In both panels, error bars depict within-subject standard error.

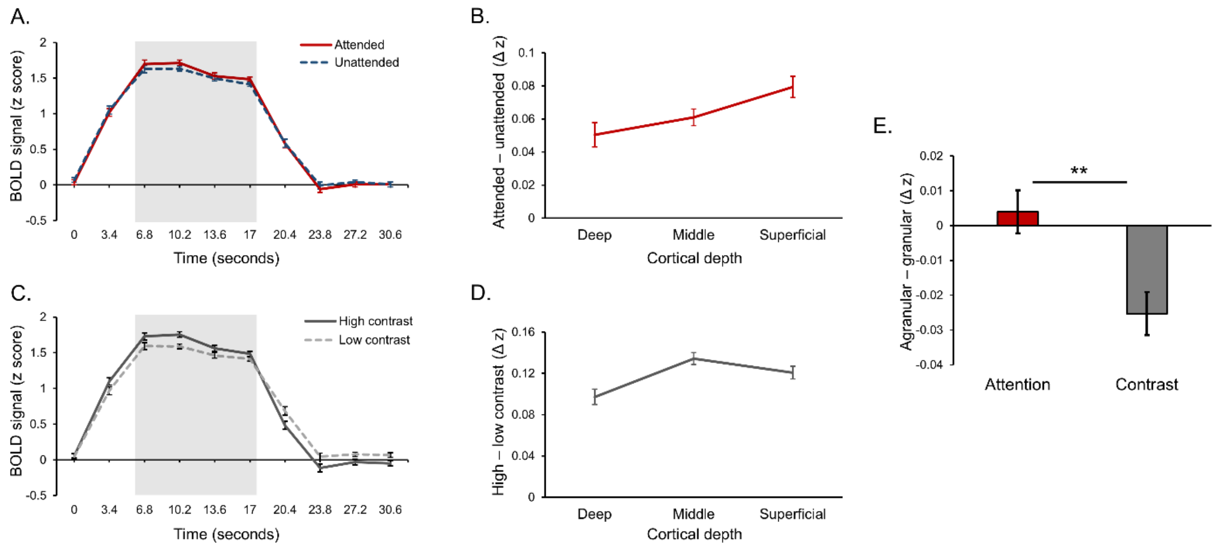


*Figure S2:* **Results using re-sampled orientation preference masks designed to sample from all cortical depths equally.** In our original analysis we selected voxels that were maximally responsive to and selective for our stimuli. This included restricting ROIs to only include voxels that exhibited a significant response to the stimuli presented in the stimulus localizer. Due to stronger overall signal in superficial cortex, this restriction created a sampling bias where our masks included more voxels that maximally overlapped with the superficial depth bin compared to other bins: on average in V1, 334 voxels (SD = 97) maximally overlapped with the superficial bin, compared to 141 (SD = 43) for middle and 175 (SD = 55) for deep. Similarly, for V2 there were 331 voxels (SD = 117) for superficial, 146 (SD = 50) for middle and 189 for deep (SD = 56) and in V3 there were 379 voxels (SD = 132) for superficial, 162 (SD = 55) for middle and 177 (SD = 54) for deep. Note that the total number of voxels reported here for each visual area do not add up to the total analyzed for each visual area (1000). This is because there were also a number of voxels that fell within white matter and CSF, which helped the spatial GLM estimate responses for these depth bins that fell outside the gray matter, but responses from these depths were not analyzed further. To check our results were not dependent on this sampling bias, we conducted the following control analysis. We resampled our orientation preference masks by randomly removing voxels until there was an equal number that maximally overlapped with each gray matter depth bin. This resulted in 132 voxels (SD = 40) for each depth bin in V1, 137 (SD = 46) for each depth in V2, and 141 (SD = 45) for each depth in V3. We recomputed our analyses using these control masks that sampled evenly from all cortical depths, which revealed very similar results. The effect of attention was significant during the highlighted time points (stats, panel **A**). The effect of attention varied across depth (F [46, 2] = 3.55, $p$ = .037, panel **B**), being larger in superficial compared to middle (t [23] = 2.05, $p$ = .052) and deep cortex (t [23] = 2.25, $p$ = .034). The effect of stimulus contrast was significant in the highlighted time window (stats, panel **C**). The effect of contrast varied significantly across depth (F [46, 2] = 5.73, $p$ = .006), being larger in the middle depth bin compared to deep (t [23] = 3.20, $p$ = .004, panel **D**). The effect of attention was significantly stronger in the agranular layers compared to stimulus contrast (t [23] = 2.37, $p$ = .026, panel **E**). Overall, the results of this control analysis were similar to our main analysis. All error bars depict within-subjects standard error.

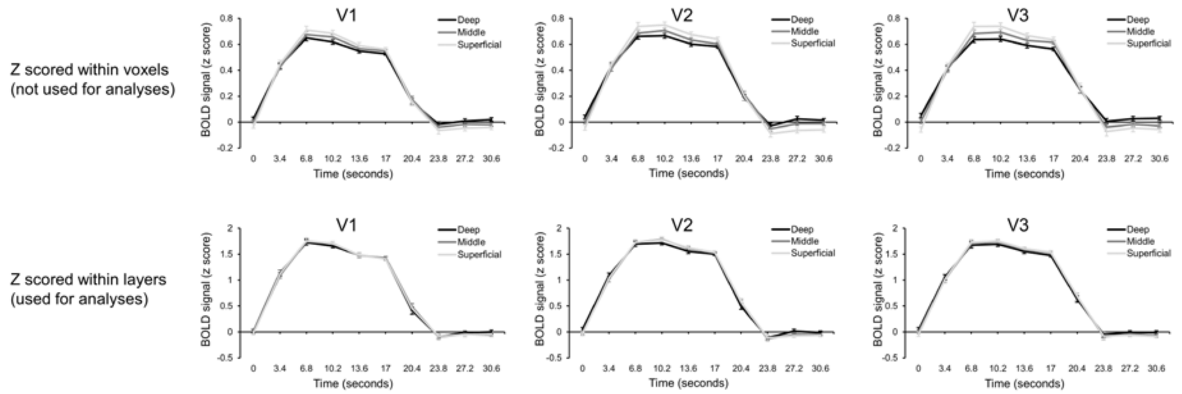


*Figure S3:* **Removal of layer-specific BOLD bias.** As expected, there was a bias in overall BOLD amplitude towards superficial cortex. This is demonstrated in the top row, which shows the average overall BOLD signal change for a block of stimuli. The data plotted in the top row were z scored within voxels, before we estimated layer-specific responses. For the analyses reported in our main paper, we removed this bias by instead z scoring data within layers, after we had estimated layer-specific responses from the pre-processed (but not z scored) voxel data. This is demonstrated in the bottom row, where the bias in BOLD amplitude in superficial cortex is almost completely removed.
